# Subcellular metabolic pathway kinetics are revealed by correcting for artifactual post harvest metabolism

**DOI:** 10.1101/749515

**Authors:** Sophie Trefely, Joyce Liu, Katharina Huber, Mary T. Doan, Helen Jiang, Jay Singh, Eliana von Krusenstiern, Anna Bostwick, Peining Xu, Juliane Bogner-Strauss, Kathryn E. Wellen, Nathaniel W. Snyder

## Abstract

**OBJECTIVE:** The dynamic regulation of metabolic pathways can be monitored by stable isotope tracing. Yet, many metabolites are part of distinct processes within different subcellular compartments. Standard isotope tracing experiments relying on analyses in whole cells may not accurately reflect compartmentalized metabolic processes. Analysis of compartmentalized metabolism and the dynamic interplay between compartments can potentially be achieved by stable isotope tracing followed by subcellular fractionation. Although it is recognized that metabolism can take place during biochemical fractionation of cells, a clear understanding of how such post-harvest metabolism impacts the interpretation of subcellular isotope tracing data and methods to correct for this are lacking. We set out to directly assess artifactual metabolism, enabling us to develop and test strategies to correct for it. We apply these techniques to examine the compartment-specific metabolic kinetics of ^13^C-labeled substrates targeting central metabolic pathways.

**METHODS:** We designed a stable isotope tracing strategy to interrogate post-harvest metabolic activity during subcellular fractionation using liquid chromatography-mass spectrometry (LC-MS).

**RESULTS:** We show that post-harvest metabolic activity occurs rapidly (within seconds) upon cell harvest. With further characterization we reveal that this post-harvest metabolism is enzymatic, and reflects the metabolic capacity of the sub-cellular compartment analyzed; but is limited in the extent of its propagation into downstream metabolites in metabolic pathways. We also propose and test a post-labeling strategy to assess the amount of post-harvest metabolism occurring in an experiment and then to adjust data to account for this. We validate this approach for both mitochondrial and cytosolic metabolic analyses.

**CONCLUSIONS:** Our data indicate that isotope tracing coupled with sub-cellular fractionation can reveal distinct and dynamic metabolic features of cellular compartments, and that confidence in such data can be improved by applying a post-labeling correction strategy. We examine compartmentalized metabolism of acetate and glutamine and show that acetyl-CoA is turned over rapidly in the cytosol and acts as a pacemaker of anabolic metabolism in this compartment.

## 1. Introduction

Metabolites function as fuel for energy production, as substrates for biosynthetic processes, and also as important signaling molecules [1,2] but the function of metabolites is highly dependent upon their sub-cellular location. For example, the central metabolite acetyl-CoA is connected to anabolic functions in the cytosol and to catabolic processes in the mitochondria. In addition, several other metabolites associated with the mitochondrial TCA cycle (including succinyl-CoA, α-ketoglutarate, succinate, fumarate) also have signaling functions in different compartments [2–5]. Stable isotope tracing by liquid chromatography-mass spectrometry (LC-MS) is a powerful technique for delineating metabolic pathways. Monitoring the incorporation of isotope labeled substrates over time can be used to determine the substrate preferences and kinetic parameters of metabolic pathways under different conditions [6]. Yet, one limitation of standard isotope tracing experiments is that analyses in whole cells may not accurately reflect compartmentalized metabolic processes. The application of sophisticated analysis to data from whole cells have provided valuable information about compartmentalized metabolic processes [7–10], but these deconvolutions can only be applied to very specific pathways and rely on annotation of metabolic pathways and assumptions which may not completely reflect the biological system being studied. Metabolic tracing in isolated organelles represents a classic and important strategy to probe distinct compartments such as mitochondria [11], but such analyses do not capture the dynamic metabolism between subcellular compartments that occurs in intact cells.

Analysis of compartmentalized metabolism can potentially be achieved by stable isotope tracing followed by subcellular fractionation, but interpretation of the resulting data is complicated due to the disruption involved in fractionation processing [12]. Thus, reliable direct measurement of kinetic parameters in sub-cellular metabolite pools is an important but challenging goal. Although it is recognized that metabolism can take place during biochemical fractionation of cells, a clear understanding of how such post-harvest metabolism impacts the interpretation of subcellular isotope tracing data and methods to correct for this are lacking. We set out to directly assess artifactual metabolism, enabling us to develop and test a post-labeling correction strategy. We apply this method and validate it in defined model systems. We demonstrate the rapid turnover of acetyl-CoA in the cytosolic compartment and its co-ordinated use in downstream anabolic pathways. We also observe distinct regulation of mitochondrial and cytosolic succinyl-CoA production and demonstrate that the use of a correction strategy is crucial for interpretation of the kinetics of mitochondrial glutamine metabolism in a labeling and fractionation experiment.

## 2. Results

### 2.1 Dual labeling strategy reveals metabolic activity during fractionation

Generation of artifacts is a concern in any bioanalytical method that includes extensive sample processing and potentially unstable analytes. This is especially relevant in sub-cellular fractionation, which is a highly disruptive process that typically requires multiple steps and at least several minutes before metabolites can be stabilized in an appropriate extraction buffer. We developed a dual labeling strategy to investigate the potential for metabolism to occur during fractionation processing (**Figure 1A**). We used Stable Isotope Labeling of Essential nutrients in Cell culture (SILEC) to introduce a heavy (^13^C_3_^15^N_1_) label into the CoA backbone of acyl-CoA molecules present in all relevant compartments in cells. Labeling of >99% of CoA across all measurable acyl-CoA species was achieved through the passaging of cells in ^15^N_1_^13^C_3_-Vitamin B5 (pantothenate) for at least 9 passages as previously described [13]. In a separate labeling scheme, we performed carbon labeling of the acyl groups of acyl-CoA molecules by incubation in ^15^N_2_^13^C_5_-glutamine for 120 min (**Figure 1A**). Glutamine metabolism results in incorporation of 4 acyl carbons into succinyl-CoA (succinyl-CoA M4) and 2 acyl carbons of acetyl-CoA (acetyl-CoA M2). The differently labeled cells (SILEC + acyl-group labeled) were combined after scraping and subjected to sub-cellular fractionation (**Figure 1A**). Thus, doubly labeled molecules with both heavy acyl and CoA groups indicate artifactual metabolism as these molecules can only form post-harvest, after the differently labeled cells have been combined (**Figure 1B, Supp. 1**). This may reflect enzyme catalyzed metabolism or non-enzymatic acyl interchange between acyl-CoA molecules [14]. Doubly and singly labeled molecules were detected by liquid chromatography-high resolution mass spectrometry (LC-HRMS) to differentiate each labeling scheme[15].

**Figure 1:**
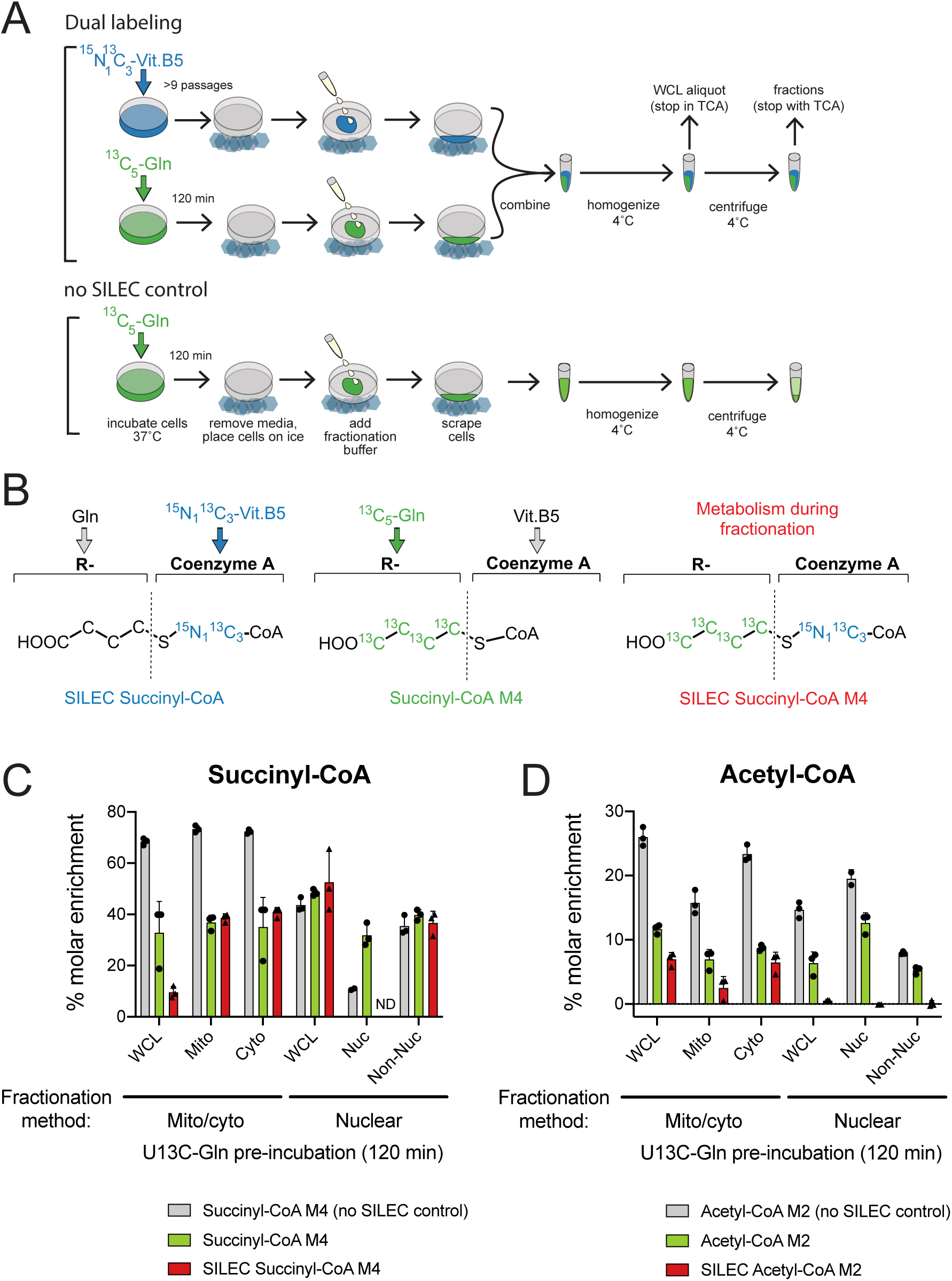
Dual labeling strategy reveals metabolism during fractionation. **A**) Schematic overview of dual labeling scheme. HepG2 cells were labeled either by culturing in ^15^N_1_ ^13^C_3_-vitamin B5 for >9 passages or incubation in DMEM containing ^15^N_2_ ^13^C_5_-glutamine (2 mM) and unlabeled glucose (5 mM) without serum for 120 min. Differently labeled cells were combined after scraping and subjected to sub-cellular fractionation with whole cell lysate (WCL) aliquot removed after homogenization. **B**) ^15^N_1_ ^13^C_5_-vitamin B5 is incorporated into the Coenzyme A backbone of acyl-CoA mole-cules (e.g. SILEC succinyl-CoA) whilst ^15^N_2_ ^13^C_5_-glutamine labels acyl (R-group) carbons (e.g. Succinyl-CoA M4). Generation of doubly labeled molecules (e.g. SILEC succinyl-CoA M4) after the labeled cells are combined indicates post-harvest metabolism during fractionation. **C, D**) Total acyl labeling was calculated for control cells that were not combined with SILEC cells as a % of the total non-SILEC acyl-CoA pool (grey bars). For dual labeling samples, the molar enrichment of acyl labelling in molecules without SILEC CoA labeling (green bars) or molecules with SILEC CoA labeling (red bars) was calculated by normalization to control samples without acyl carbon labeling i.e. ^15^N_2_ ^13^C_5_-glutamine was replaced with unlabeled glutamine. n=3 distinct replicate samples, error bars indicate standard deviation.

When applied to two classical differential centrifugation-based fractionation methods (mitochondria/cytosolic and nuclear (**Supp. 2B**)), the dual labeling strategy revealed extensive artifactual metabolism during sub-cellular fractionation (red bars). The extent of artifactual post-harvest metabolism observed varied across different fractionation methods, subcellular fractions, and metabolites analyzed. Notably, formation of doubly labeled succinyl-CoA from glutamine was generally more extensive than for acetyl-CoA, proportional to acyl labeling observed in the control with only ^15^N_2_^13^C_5_-glutamine labeled cells (**Figure 1C,D**). For both acetyl-CoA and succinyl-CoA, the extent of artifactual metabolism detected in the nucleus was markedly less than in the mitochondria and cytosol (**Figure 1C,D**).

### 2.2 Substrate use during fractionation follows defined metabolic pathways

Having established that artifactual metabolism occurs during sub-cellular fractionation, we conducted a series of experiments to further evaluate the artifact’s nature and extent. We designed an experimental approach to trace artifactual metabolism by addition of ^13^C-labeled substrates to the fractionation buffer (**Figure 2A**). The detection of ^13^C label in metabolites of interest thus reflected metabolic activity during processing. The introduction of ^13^C-substrates from the very first step of cell processing (on cold cells before they were scraped) allowed us to monitor the extent of artifactual metabolism over the entire procedure.

**Figure 2:**
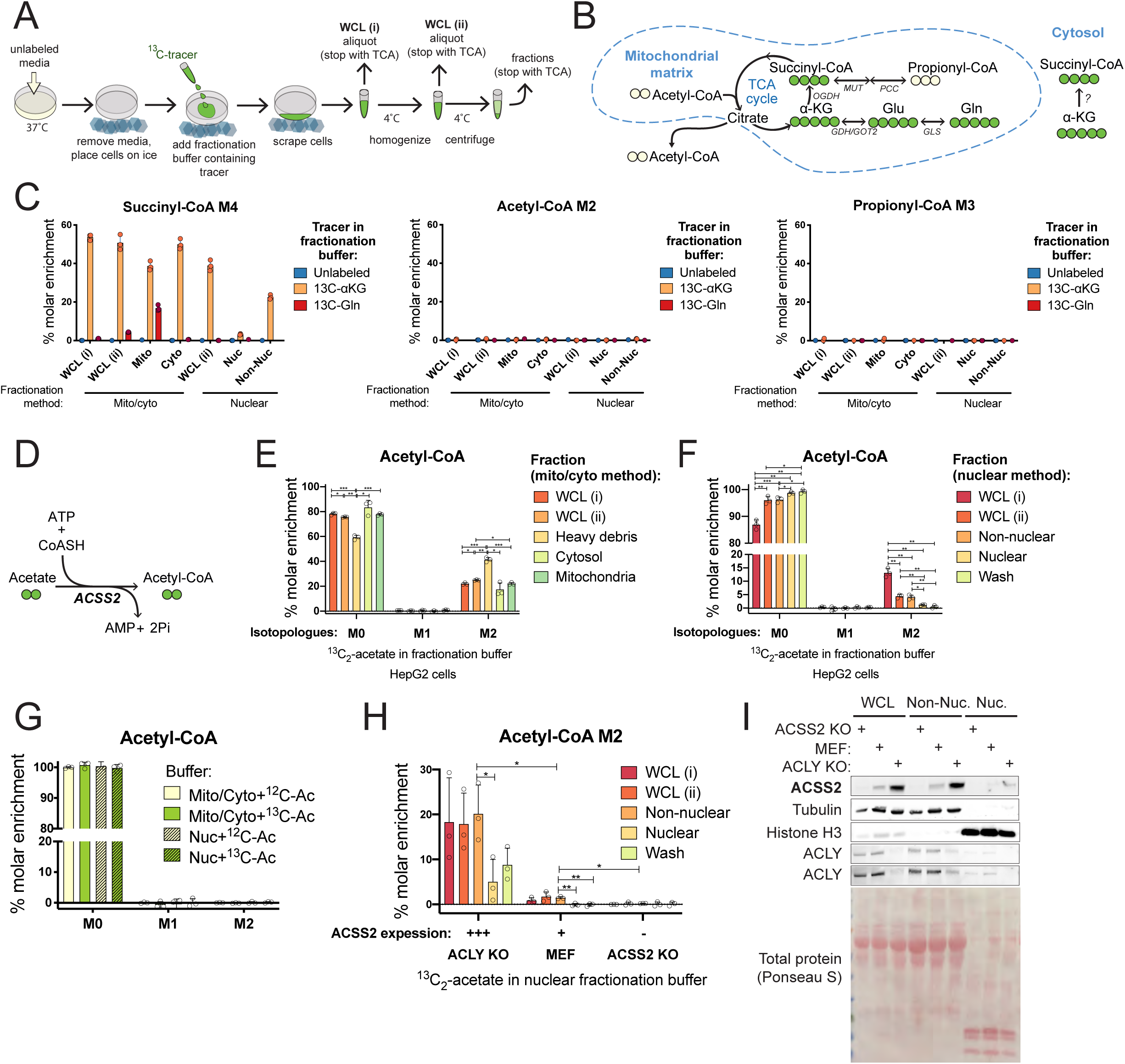
Substrate use during fractionation follows defined metabolic pathways. **A**) Schematic overview of labeling approach to test for post-harvest metabolism. Cells were cultured in unlabeled media and ^13^C-labeled substrate was introduced in the fractionation buffer before fractionation processing and quenching of fractionations in trichloroacetic acid (TCA). **B**) Incorporation of ^15^N_2_ ^13^C_5_-glutamine or ^13^C_5_ -α-ketoglutarate into the mitochondrial TCA cycle results in labelling of 4 acyl carbons of succinyl-CoA. The enzymatic pathway for incorporation of ^13^C_5_ -α-ketoglutarate into succinyl-CoA M4 in the cytosol is not defined. **C**) Hepatocellular carcinoma (HepG2) cells were harvested in fractionation buffer containing either 0.5 mM ^15^N_2_ ^13^C_5_-glutamine or 0.5 mM ^13^C_5_ -α-ketoglutarate and % molar enrichment of acyl labelling was calculated by normalization to unlabeled control samples **D**) Acyl-CoA synthetase short-chain family member 2 (ACSS2) catalyzes the conversion of acetate to acetyl-CoA. HepG2 cells were harvested in fractionation buffer containing 0.1 mM ^13^C_2_-acetate and % molar enrichment of acetyl-CoA isotopologues was calculated by normalization to control samples with unlabeled acetate in the fractionation buffer for mito/cyto (**E**) and nuclear (**F**) fractionation protocols. **G**) Isotopologue enrichment was analyzed in de-proteinated HepG2 cell extracts resuspended in fractionation buffer for mito/cyto or nuclear protocols containing 0.1 mM ^13^C_2_-acetate or unlabeled acetate and incubated on ice for 1 h before extraction. **H**) ACLY KO, control and ACSS2 KO MEF cells were harvested in fractionation buffer containing 0.1 mM ^13^C_2_-acetate and acetyl-CoA M2 enrichment calculated by normalization to control samples harvested in fractionation buffer containing 0.1 mM unlabeled acetate. **I**) Western blot analysis comparing ACSS2 expression levels in ACLY KO, control and ACSS2 KO MEF cells with equal protein loading. Tubulin and histone H3 serve as loading controls and fraction-specific markers for non-nuclear and nuclear fractions, respectively. The fractionation buffers for both these methods are hypotonic sugar solutions, with the most notable difference being the presence of dilute detergent (0.1% NP-40) in the nuclear fractionation buffer. This detergent destroys the mitochondrial membrane such that the non-nuclear fraction is expected to contain soluble mitochondrial contents. Representative experiments are shown with bars representing the mean of n=3 distinct replicate samples (induvidual symbols) and error bars indicating standard deviation. For comparison between two groups, datasets were analyzed by two-tailed student’s t-test with statistical significance defined as p < 0.05 (*), p < 0.01 (**), p < 0.001 (***).

The extent of glutamine incorporation into succinyl-CoA in the post-labeling paradigm (**Figure 2C**) is much lower than that observed after 120 min pre-labeling of cells (**Figure 1C**) since during pre-incubation, carbon from glutamine will have time to accumulate in both succinyl-CoA and various other metabolic products. Therefore, to examine the artifactual activity of the pathway for conversion of glutamine to succinyl-CoA, we tested post-harvest metabolism of both ^15^N_2_^13^C_5_-glutamine and its metabolic product ^13^C_5_-α-ketoglutarate (**Figure 2B**). Post-harvest incorporation of ^13^C_5_-α-ketoglutarate into succinyl-CoA M4, was substantial in most fractions, while substantial labeling from ^13^C-glutamine was only observed in mitochondria (**Figure 2C**). Importantly, this metabolism occurs rapidly, with ^13^C_5_-α-ketoglutarate conversion to succinyl-CoA M4 representing >40% of the succinyl-CoA pool in the whole cell lysate extracted immediately after cell scraping (WCL i) in both the mito/cyto and the nuclear isolation procedure. That incorporation of ^15^N_2_^13^C_5_-glutamine in the fractionation buffer into succinyl-CoA was limited compared to ^13^C -α-ketoglutarate (**Figure 2C**) is not surprising, given that glutamine is separated from succinyl-CoA by 3 metabolic steps, and α-ketoglutarate only by 1 step (**Figure 2B**). ^13^C_5_ -α-ketoglutarate or ^15^N_2_^13^C_5_-glutamine labeling of distal metabolites, acetyl-CoA and propionyl-CoA, was not detected (**Figure 2C**). In support of this interpretation, acetate conversion to acetyl-CoA is a single step reaction and occurs efficiently with the addition of 0.1 mM ^13^C_2_-acetate to fractionation buffer, producing acetyl-CoA M2, which makes up 40% of the cytosolic pool in HepG2 cells (**Figure 2D,E,F**). ^13^C_2_-acetate added to the fractionation buffer was also detectable in 3-hydroxymethylglutaryl-CoA (HMG-CoA), a proximal metabolite in the mevalonate pathway, with predictable 2-carbon inclusion patterns signifying the co-ordinated addition of acetyl units (**Supp. 2A,C**). Acetate labeling was generally limited to low levels in downstream metabolites (**Supp. 2C**). Thus, artifactual metabolism during fractionation is limited in extent to proximal reactions in metabolic pathways.

Each sub-cellular fraction exhibits distinctive enrichment of appropriate proteins (**Supp. 2B**). To investigate the idea that compartmentalization of proteins could drive post-harvest metabolism, we tested for enzyme dependence of artifactual metabolism. We first used methanol:water (80:20) to precipitate protein from whole HepG2 cells and separate it from soluble metabolites. De-proteinated cell extracts were resuspended and incubated in fractionation buffer containing ^13^C_2_-acetate. Acetyl-CoA M2 enrichment was not detected after incubation (**Figure 2G, Supp. 2D**) indicating that artifactual acetate conversion to acetyl-CoA is protein dependent. To specifically establish the enzyme dependence of artifactual metabolism, we compared 3 different mouse embryonic fibroblast (MEF) cell lines, which allowed us to titrate the expression levels of acyl-CoA synthetase short-chain family member 2 (ACSS2), the nuclear-cytosolic enzyme that produces acetyl-CoA from acetate (**Figure 2H**). ACLY loss activates a glucose-to-acetate metabolic switch, driven by the upregulation of ACSS2 for the production of cytosolic acetyl-CoA, which is used in downstream anabolic pathways in the cytosol including lipid synthesis. Acetate accounts for ∼80 % of the total acetyl-CoA pool in these cells [16]. The high ACSS2 expression in the ACLY KO MEF cells corresponded to greater artifactual generation of acetyl-CoA M2 during fractionation. ACLY KO MEFs displayed 20% enrichment in the non-nuclear fraction (which includes the cytosol), whilst the modest ACSS2 expression in control MEF cells was accompanied by similarly modest artifactual acetyl-CoA M2 generation (1.4 %) and acetyl-CoA M2 was barely enriched (0.2 %) in ACSS2 KO MEFs (**Figure 2H**). Thus, artifactual metabolism is enzymatic and reflects the specific metabolic capacity of the cell type and compartment isolated.

Two principle strategies to avoid artifactual metabolic activity are speed and cold. Since we have shown that artifactual metabolism occurs to a large extent upon cell scraping, a process that requires less than 1 minute (**Figure 2C,E,F**) it is unlikely that any fractionation method that uses cultured cells harvested by scraping can avoid substantial artifactual metabolism regardless of the speed of the subsequent steps. To address the effect of cold, we performed tracing of ^13^C-acetate in the fractionation buffer under extreme cold conditions. Cells were scraped at −5 °C (the sucrose buffers did not freeze at this temperature). Centrifugation steps were also performed at −5 °C and all transfer steps were performed in a cold room (4 °C) with samples in −5 °C isopropanol bath. This extra cold processing, however, did not prevent artifactual conversion of ^13^C-acetate to acetyl-CoA M2 in either the mito/cyto or the nuclear isolation procedures (**Supp. 2E,F**).

### 2.3 Rapid quenching of whole cells reveals the dynamic nature of acetyl-CoA metabolism

Given that post-harvest metabolism occurs, we next sought to develop a strategy account and correct for this. We first targeted acetate and acetyl-CoA metabolism specifically in the cytosol. We again took advantage of the avid acetate usage in ACLY KO MEF cells. Notably, in these cells, exogenous acetate feeds cytosolic acetyl-CoA pools as well as fatty acid synthesis, but is not accompanied by increased mitochondrial acetate usage. Tracing 0.1 mM ^13^C-acetate into intermediates of the TCA cycle revealed negligible incorporation in ACLY KO MEFs [16]. Thus, whole cell acetate tracing in ACLY KO MEFs largely represents the cytosolic acetate utilization, allowing us to interrogate the dynamics of the cytosolic acetyl-CoA pool via direct extraction of metabolites from whole cells (i.e., a ‘pseudo-cytosolic’ system). This metabolic wiring (**Figure 3A** makes these cells uniquely useful for validating results obtained from sub-cellular tracing by allowing us to compare whole cell ‘pseudo-cytosolic’ data, obtained immediately upon cell harvest, with cytosolic data obtained through sub-cellular fractionation.

**Figure 3:**
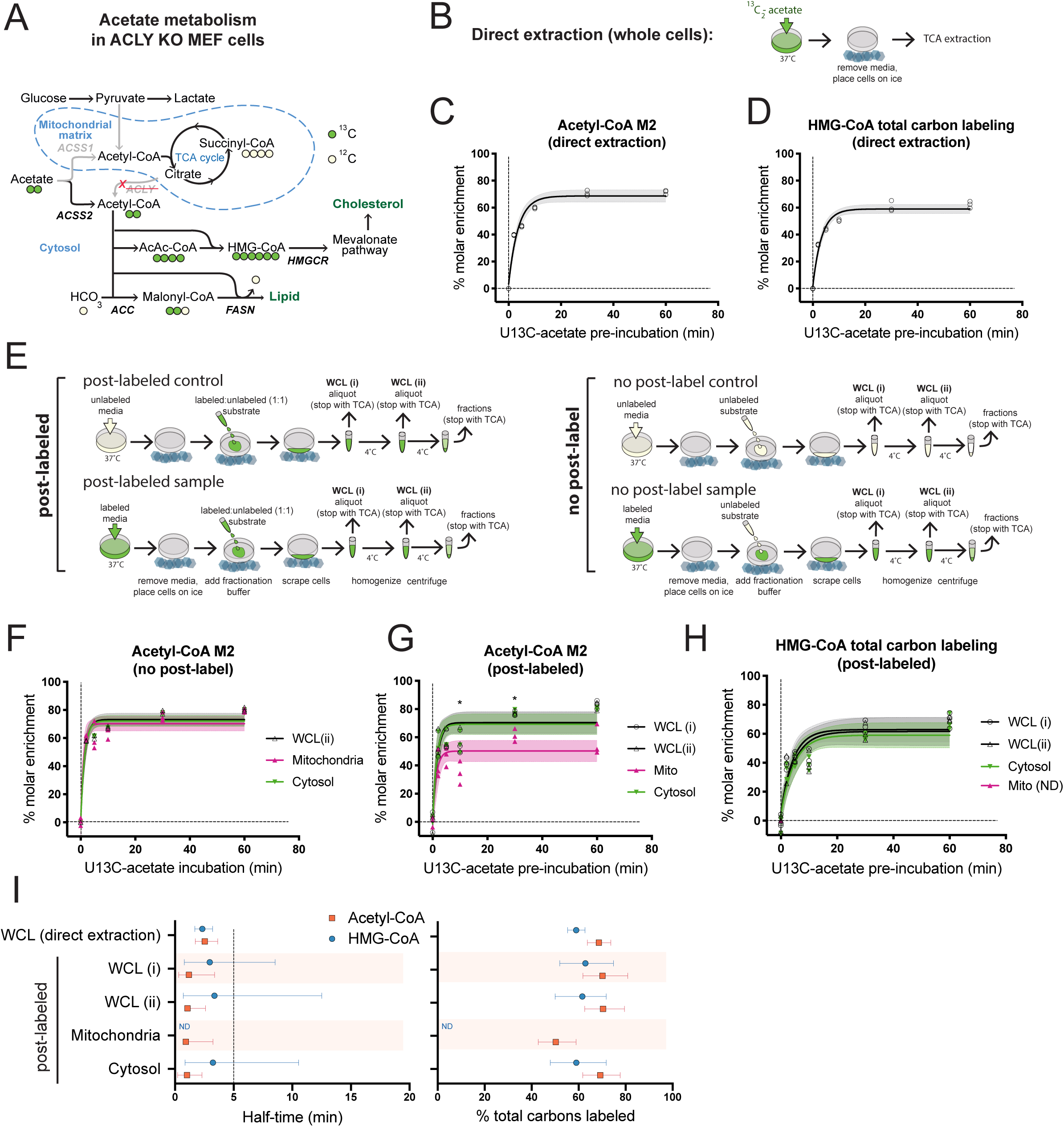
Post-labeling correction resolves mitochondrial and cytosolic acetyl-CoA metabolism. ACLY KO MEF cells were pre-labeled by incubation with 0.1 mM ^13^C_2_-acetate over a time-course. **A**) Acetate is preferentially used for cytosolic acetyl-CoA generation in ACLY KO MEFs. **B**) In direct extraction, whole cells were immediately quenched in trichloroacetic acid (TCA). **C**) Acetyl-CoA M2 enrichment after direct extraction. **D**)3-hydroxymethylglutaryl-CoA (HMG-CoA) total acyl carbon labeling, indicating % enrichment of all 6 acyl carbon atoms after direct extraction. **E**) Schematic comparison of post-labeled and no post-label approach. Post-labeling accounts for artifactual metabolism during processing by addition of a supraphysiological concentration of partially labeled tracer to cells that were pre-labeled with tracer as well as controls that were not pre-labeled. No tracer is added upon cell harvest in the no post-label approach. **F**) Representative data from time-course for enrichment of acetyl-CoA M2 into whole cell lysate (WCL), mitochondria and cytosol after sub-cellular fractionation with no post-labeling. Enrichment was calculated by normalization to unlabeled controls (time=0). **G**) Representative data from time-course for enrichment of acetyl-CoA M2 and (**H**) HMG-CoA in whole cell lysate (WCL), mitochondria and cytosol after sub-cellular fractionation with post-labeling by inclusion of 5 mM each of ^13^C_2_-acetate and unlabeled acetate in the fractionation buffer. Enrichment was calculated by normalization to post-labeled controls (time=0). **I**) Comparison of direct extraction from whole cells and post-label normalization of sub-cellular fractions at steady-state. Representative experiments with n=3 distinct replicate samples (induvidual symbols) are shown and error bars indicating 95% confidence bands. For **C,D,F**-**H**, shading indicates 95% confidence bands and symbols represent individual replicate values. For **F** and **G**, statistical significance was determined for comparison between mitochondrial and cytosolic data for each time point by two-tailed student’s t-test analysis with significance defined as p < 0.05 (*). ND = not determined.

We validated a robust technique for harvesting whole cells in stable isotope tracing experiments within this system by comparing ^13^C_6_-glucose tracing into acetyl-CoA in ACLY KO MEF cells using 4 different extraction methods head to head. Immediately quenching the metabolism of live cells with cold solvent produced consistent results - direct extraction into cold (4 °C) 10% (w/v) trichloroacetic acid in water (TCA) and direct extraction into cold (−80 °C) methanol:water (80:20) yielded identical results (5.4% acetyl-CoA M2) showing low levels of glucose incorporation consistent with previous data [16]. Modifications from direct extraction with additional PBS washing or centrifugation steps, however, produced disparate results (**Supp. 3A**).

Having established a gold standard result for the whole cell labeling by direct extraction into 4 °C 10% TCA, we applied this to assess ‘pseudo-cytosolic’ acetyl-CoA turnover. ACLY KO MEFs were incubated in physiological (0.1 mM) acetate concentration and ^13^C enrichment of acetyl-CoA and a panel of short chain acyl-CoA metabolites was monitored over time upon switching to ^13^C_2_-acetate media. Acetate was rapidly incorporated into the acetyl-CoA pool, with a steady-state acetyl-CoA M2 level of 69 % achieved in a half-time of less than 5 min (**Figure 3B,H)** The incorporation of carbons from acetate into acetyl-CoA (acetyl-CoA M2) is rapidly propagated into downstream metabolites including the mevalonate pathway metabolite HMG-CoA (**Figure 3C,H; Supp. 3I**). These data were recapitulated in a complementary, reverse labeling approach, monitoring loss of ^13^C-label in cells that were pre-labeled for 16 h with ^13^C-acetate, before switching to media containing unlabeled acetate (**Supp. 3G,I**).

### 2.4 Post-labeling correction resolves mitochondrial and cytosolic acetyl-CoA metabolism

To develop a strategy to account for post-harvest metabolism, we hypothesized that introduction of an overwhelming supraphysiological concentration of a 1:1 labeled:unlabeled mixture of the experimentally traced substrate to the fractionation buffer of cells that had been incubated in the absence of labeled substrate could be used to generate a control for data correction. The overwhelming concentration is intended to force artifactual metabolism in the same direction by mass action with a 1:1 chance of incorporating either no label or the full label (preventing change or saturation of enrichment) regardless of different metabolic conditions across experimental groups. To study the metabolism of experimental groups, cells prelabeled with the substrate were processed in parallel with the same partially labeled buffer (**Figure 3D**). The substrate used in the fractionation buffer was identical to the test substrate to reflect the targeted metabolic pathways. An additional control with no label was used to assess the absolute degree of labeling.

^13^C_2_-acetate tracing followed by isolation of the mitochondrial and cytosolic fractions was performed with and without post-labeling correction and compared to the ‘pseudo-cytosolic’ data. For post-labeling an overwhelming concentration of 1:1 labeled:unlabeled acetate (5 mM each) was added to the fractionation buffer. We are not aware of reliable data on the intracellular concentration of free acetate, however, the concentration of acetate in human plasma is 50 – 250 µM [17] and although it is possible that accumulation of acetate within cells may occur, the observation of dynamic acetate recycling and efflux suggests that this is not likely [18]. Thus the combined post-labeling concentration of 10 mM is likely ∼50-fold the cellular acetate concentration, which is diluted in fractionation buffer upon cell homogenization. Strikingly, post-label correction (i.e. post-labeled experimental groups normalized to post-labeled controls) reveals distinct labeling patterns for acetyl-CoA M2 enrichment in the mitochondria and the cytosol whereas no post-labeling (i.e. no post-label experimental groups normalized to no post-label controls) does not (**Figure 3F,G**). Lower steady-state acetate labeling in the mitochondria is consistent with the flexibility of mitochondrial acetyl-CoA pools, which can draw from a variety of alternative substrates including glucose, whilst cytosolic acetyl-CoA sources are more limited and acetate dependent in the ACLY KO MEF model (**Figure 3E**). Notably, incorporation of acetate into the mitochondrial TCA cycle intermediate succinyl-CoA was ∼1% (**Supp. 3E,F**), supporting minimal mitochondrial acetate use in ACLY KO MEF cells. Comparison of post-labeling correction data to the ‘pseudo-cytosolic’ data for acetyl-CoA showed that acetate kinetics in isolated cytosol and in whole cells with rapid quenching closely agree (**Figure 3B,G,H Supp. 3I**), consistent with the majority of the acetyl-CoA pool being localized to the cytosol in these cells. The consistency of these data strongly supports the use of post-labeling as an effective normalization strategy since artifactual labeling of acetyl-CoA during post-labeling is as high as 20% (**Supp. 3D**). Mitochondrial HMG-CoA can appear as an intermediate in the ketogenesis and leucine catabolism pathways, however HMG-CoA was barely, if at all detected in the mitochondrial fraction, indicating that in ACLY KO MEFs under these conditions the dominant HMG-CoA pool is present in the cytosol where it contributes to the mevalonate pathway. Accordingly, incorporation of acetate into cytosolic HMG-CoA is consistent with ‘pseudo-cytosolic’ direct extraction (**Figure 3C,H,I; Supp. 3I**). The half-time and steady-state level of acetate incorporation into cytosolic acetyl-CoA and downstream into cytosolic HMG-CoA are consistent in both ‘pseudo-cytosolic’ and cytosolic fractionation experiments with post-label correction (**Figure 3H**). The synchronization of label incorporation in these pathways demonstrates the rapid transferal of cytosolic acetyl-CoA into downstream anabolic pathways.

### 2.5 Kinetics of mitochondrial glutamine metabolism are transformed by post-label correction

To further test the generalizability of our post-labeling approach we applied it to examine the kinetics of succinyl-CoA metabolism by incubation of HepG2 cells in ^15^N_2_,^13^C_5_-glutamine over a time-course before analysis of isotopologue enrichment into short chain acyl-CoAs in the mitochondria and cytosol. The kinetics of succinyl-CoA M4 labeling were strikingly different between compartments (**Figure 4A**). Mitochondrial glutamine incorporation appeared to be very rapid (half-time of 4.1 min) compared to the cytosol (half-time of 14.4 min), though the percent molar enrichment of succinyl-CoA M4 in both pools converge at a steady state to 76 % labeling after 80 min ^15^N_2_^13^C_5_-glutamine incubation. The clear separation at early timepoints of succinyl-CoA labeling between mitochondrial and cytosolic pools indicates that the mitochondria remain intact and mostly impermeable to succinyl-CoA exchange during fractionation, since the 2 compartments are only physically separated at the final step of processing. Acetyl-CoA and propionyl-CoA also exhibited distinct kinetics of labeling in mitochondria as compared to cytosol (**Supp 4A,B**). The much faster labeling of mitochondrial than whole cell succinyl-CoA is inconsistent with the presumption that much of cellular succinyl-CoA is contained within mitochondria. We therefore questioned how much of this mito/cyto labeling was due to metabolism that took place during the intended ^15^N_2_,^13^C_5_-glutamine incubation period versus artifactual post-harvest metabolism during the fractionation process. Since post-harvest conversion of ^15^N_2_, ^13^C_5_-glutamine into succinyl-CoA M4 occurred to a greater extent in the mitochondria than the cytosol and WCL (**Figure 2C**) this could conceivably lead to apparent (but artifactual) kinetic differences observed between these compartments in cells pre-incubated in ^15^N_2_, ^13^C_5_-glutamine (**Figure 4A**).

**Figure 4:**
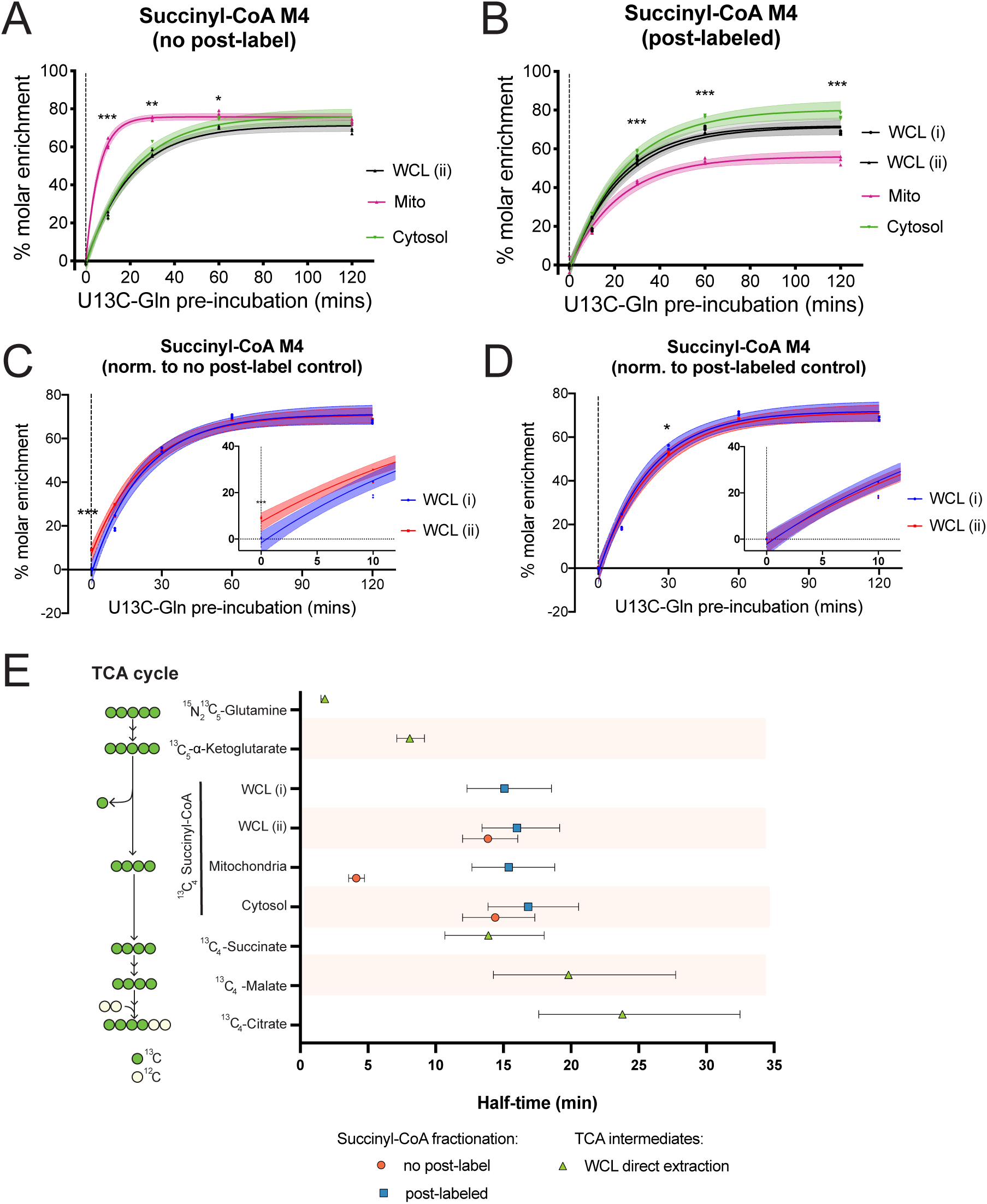
Kinetics of glutamine metabolism in the mitochondria are transformed by post-label correction. HepG2 cells were pre-labeled over a time-course with 2 mM ^15^N_2_ ^13^C_5_-glutamine. **A**) Representative data from time-course for enrichment of succinyl-CoA M4 into whole cell lysate (WCL), mitochondria and cytosol in ACLY KO MEF cells after sub-cellular fractionation in the absence of post-labeling. Enrichment was calculated by normalization to unlabeled controls (time=0) **B**) Representative data from time-course for enrichment of succinyl-CoA M4 in whole cell lysate (WCL), mitochondria and cytosol after sub-cellular fractionation with post-labeling by inclusion of 5 mM each of ^15^N_2_ ^13^C_5_-glutamine and unlabeled glutamine in the fractionation buffer. Enrichment was calculated by normalization to post-labeled controls (time=0). **C**) Comparison of absolute labeling in WCL i and WCLI ii post-labeled samples by normalization to no label control. **D**) Comparison of corrected WCL i and WCLI ii post-labeled data when normalized to post-label control. **E**) Comparison of half-times to achieve steady-state labeling for tricarboxylic acid (TCA) cycle intermediates after direct extraction from whole cells, and sub-cellular fractionation data with and without post-labeling correction. Mean of n=3 distinct replicate samples from representative experiments is shown with error bars indicating 95% confidence bands. For **A-D**, shading indicates 95% confidence bands and symbols represent individual replicate values. For comparison between two groups at each time point, datasets were analyzed by student’s t-test with statistical significance defined as p < 0.05 (*), p < 0.01 (**), p < 0.001 (***). For **A** and **B** statistical difference between mitochondrial and cytosolic data for each time point is displayed.

We investigated the impact of post-harvest metabolism on succinyl-CoA labeling kinetics using our post-labeling approach. HepG2 cells were preincubated in ^15^N_2_, ^13^C_5_-glutamine over a time-course followed by processing with 1:1 ^15^N_2_, ^13^C_5_-glutamine:glutamine (both at 5 mM). Intracellular glutamine concentrations between 4 µM and 1.4 mM have been reported [19]. Thus the combined post-labeling concentration of 10 mM is likely >10-fold the cellular glutamine concentration, and cell contents are significantly diluted upon addition of fractionation buffer. Post-label normalization led to a substantial transformation of the mitochondrial succinyl-CoA labeling, whilst having little impact on the cytosolic and WCL data (**Figure 4B**). Several factors indicate the validity of the post-label corrected data. First, the WCL now agrees with the average of cytosol and mitochondria, consistent with at least a substantial proportion of the actively metabolized succinyl-CoA pool being localized to the mitochondria. Second, when the data are normalized to post-label controls the WCL (i) and WCL (ii) curves overlap (**Figure 4C**). Normalization of the post-labeled data for WCL to no post-label controls reveals the absolute level of labeling is lower in the WCL(i) aliquot collected earlier in processing compared to WCL(ii), collected after cell homogenization (**Figure 4D**). Since WCL(i) is the same fraction as WCL(ii) except with less artifactual metabolism, the data should converge, as observed, if artifactual metabolism is corrected for. Third, analysis of ^15^N_2_,^13^C_5_-glutamine incorporation into other TCA cycle intermediates directly extracted from whole cells show half-times consistent with the post-labeled corrected data and inconsistent with uncorrected data (**Figure 4E, Supp 4F-J**). Thus, we conclude that inclusion of the post-label correction prevents erroneous conclusions and reveals the true kinetics of mitochondrial glutamine metabolism.

## 3. Discussion

In sum, the data clearly demonstrate that metabolism occurs post-harvest, that it cannot be sufficiently prevented by rapid speed of fractionation or by very cold temperatures, and that this impacts the interpretation of data from experiments involving isotope tracing followed by subcellular fractionation. Kinetic data in mitochondrial fractions appear to be particularly compromised by post-harvest artifactual metabolism, at least during glutamine tracing. We further show that a post-label correction can be applied to improve confidence in data from such experiments. As interest in the role of metabolites as signaling molecules and the importance of compartmentalization in their function gains pace, sub-cellular metabolomic analyses are emerging as a critical tool [19–22]. Though we have used classical fractionation approaches in this study, the post-label correction strategy can be applied to any fractionation approach in which the investigator employs metabolic labeling followed by fractionation. Our strategy presents a useful approach to help validate previous and emerging sub-cellular tracing studies in which compartments are isolated after experimental perturbation [23]. This is a powerful approach as it provides insight into the dynamic metabolic interplay between organelles.

Several important caveats limit the applicability of post-labeling as a universal correction factor. We demonstrate that multiple proximal metabolic intermediates contribute to artifactually generated pools of metabolites (**Figure 2**), thus the validity of just one tracer as a correction factor is not comprehensive. Practical limitations make the addition of tracers accounting for all metabolic steps between a tracer and the measured metabolite unfeasible, especially in cases where the exact metabolic pathway generating a metabolite in a specific compartment may not be known. Moreover, access to the correction tracer in discrete membrane bound compartments such as the mitochondria may be limited by the availability and activity of transporters. The post-labeling approach also by definition can only be applied to tracing experiments. While post-harvest metabolism may also impact interpretation of subcellular metabolite quantification, different strategies will be required to address non-isotope tracing experiments.

Despite these caveats and limitations, our approach represents an important step towards resolving metabolic fluxes in subcellular compartments. We directly demonstrate the validity of post-labeling correction for the analysis of glutamine and acetate metabolism in the mitochondria and cytosol and reveal kinetic data with direct physiological relevance. We show rapid acetyl-CoA turnover specifically in the cytosol and its co-ordinated transferral downstream into the mevalonate pathway, implicating cytosolic acetyl-CoA as a dynamic pacemaker for anabolism. We also reveal distinct regulation of mitochondrial from cytosolic succinyl-CoA - a metabolite pool whose existence is indicated by succinylation of cytosolic proteins [24,25], yet whose metabolic origin and regulation is not defined. We anticipate that this post-labeling strategy can be applied to address numerous biological questions about compartmentalized metabolic regulation.

## 4. Experimental Procedures

### 4.1 Cell culture

Cells were passaged every 2-3 days at 80% confluence. Mouse embryonic fibroblast (MEF) cell lines were generated in the Wellen lab[16] (available upon request) and were cultured in DMEM, high glucose (Thermo Fisher Scientific, Gibco #11965084) with 10% calf serum (Gemini bio-products #100-510, lot C93GOOH). Hepatocellular carcinoma (HepG2) cells were used at <20 passages from ATCC stocks (ATCC cat. #HB-8065) and were cultured in DMEM, high glucose with 10% fetal bovine serum (Gemini Biosciences) and penicillin/streptomycin (Thermo Fisher Scientific, Gibco cat. #10378016). All cells were tested and mycoplasma-free.

### 4.2 Stable isotope pre-labeling

For glutamine tracing, normal growth media was aspirated and replaced with unlabeled DMEM (5 mM Glc, 2 mM Gln, no serum) and cells were preincubated for 1 h. The pre-incubation media was replaced with ^13^C-Gln media (DMEM without glutamine, glucose or phenol red (Thermo-Fisher scientific, Gibco cat. #A1443001) supplemented with 5 mM glucose and 2 mM ^5^N_2_ ^13^C_5_-glutamine added in a backwards time-course, starting with 2 h, so that all dishes finish at the same time.

For acetate tracing, normal growth media was aspirated and replaced with unlabeled DMEM + 0.1 mM acetate (high glucose DMEM + 10% dialyzed FBS and 0.1 mM unlabeled acetate) and cells were preincubated for 1 h. The pre-incubation media was replaced with ^13^C-acetate media (high glucose DMEM + 10% dialyzed FBS and 0.1 mM ^13^C_2_-acetate added in a backwards time-course, so that all dishes finish at the same time.

SILEC labeling of HepG2 cells was achieved as previously described [26]-HepG2 cells were passaged in custom DMEM (high glucose (25 mM), glutamine (2 mM)) without pantothenate (Athena Biosciences cat. #0500-198) with the addition of ^15^N_1_ ^13^C_3_-vitamin B5 (pantothenate) (Isosciences) and charcoal:dextran stripped fetal bovine serum (Gemini Biosciences cat. #100-199, lot #A67FO2H) to 10%. Labeling efficiency was tested by isotopologue enrichment analysis with comparison to unlabeled controls cells.

For glucose labeling of ACLY KO MEF cells, normal growth media was aspirated and replaced with unlabeled DMEM (hi glucose DMEM + 10% dialyzed FBS + 1 mM acetate) and cells were preincubated for 16 h. The pre-incubation media was replaced with ^13^C_6_-glucose media (DMEM without glucose or glutamine (Gibco) supplemented with 2 mM glutamine and 5 mM ^13^C_6_-glucose + 10% dialyzed FBS, no acetate) for 4 h. Control unlabeled cells received the same media except with unlabeled glucose instead of ^13^C_6_-glucose.

### 4.3 Mitochondria and cytosol isolation

Media was poured from dishes into a waste container and cells were placed on ice at a 45° angle and residual media drained and aspirated completely. Dishes were laid flat on ice and 1 ml ice-cold buffer (210 mM mannitol, 70 mM sucrose, 5 mM Tris-HCl (pH 7.5), 1 mM EDTA (pH 8), adjusted to pH 7.5) added to each dish. Cells were scraped into the buffer and transferred to a pre-chilled 1 ml Potter-Elvehjem Tissue Grinder (Corning cat. #7725T-1) in a beaker of ice and water. For WCL i analysis, 100 µl was removed after scraping and quenched in 1 ml ice-cold 10% trichloroacetic acid (TCA) (Sigma cat. #T6399) in deionized water. Cells were lysed by stroking with the pestle operated at 1,600 rpm (15 strokes for HepG2 cells, 30 strokes for MEF cells). Homogenate was transferred to 1.5 ml tubes on ice. For WCL ii analysis, a 100 µl aliquot of homogenate was removed and quenched in 1 ml ice-cold 10% TCA in water. Homogenate was centrifuged at 1,300 × g from 10 min at 4 °C and supernatant was transferred to a new pre-chilled 1.5 ml tube. The ‘heavy debris’ pellet was quenched by resuspension in 1 ml 10% TCA and the supernatant was centrifuged at 10,000 × g for 20 min at 4 °C to pellet mitochondria. The supernatant (the cytosolic fraction) was quenched by transferal to a new 1.5 ml tube containing 0.25 ml of 50% (w/v) TCA in water to make a final concentration of 10% TCA. Residual cytosolic fraction was removed from the mitochondrial pellet with P200 pipette, and the pellet was quenched by resuspension in 1 ml 10% w/v TCA in water. Samples were stored at-80 before thawing for processing or directly processed.

### 4.4 Nuclear isolation

Media was poured from dishes into a waste container and cells were placed on ice at a 45° angle and residual media drained and aspirated completely. Dishes were laid flat on ice and 0.5 ml ice-cold lysis buffer (250 mM sucrose, 15 mM Tris-HCl (pH 7.5), 60 mM KCl, 15 mM NaCl, 5 mM MgCl_2_, 1 mM CaCl_2_ and NP-40 0.1%, adjusted to pH 7.4) added to each dish. Cells were scraped into the buffer. For WCL i analysis, 50 µl was removed after scraping and quenched in 1 ml ice-cold 10% TCA in water. Cell suspension was homogenized by pipetting up and down on the dish with a P1000 4 times and transferred to a pre-chilled 1.5 ml tube on ice. For WCL ii analysis, 50 ul was removed after homogenization and quenched in 1 ml ice-cold 10% TCA in water. Nuclei were pelleted by centrifugation at 600 × g for 5 min at 4 °C. The supernatant (the ‘non-nuclear’ fraction) was quenched by transferal to a new 1.5 ml tube containing 0.125 ml of 50% (w/v) TCA in water to make a final concentration of 10% TCA. The nuclear pellet was washed by the addition of 0.5 ml lysis buffer without NP-40 and recentrifuged at 600 × g for 5 min at 4 °C. For experiments in which tracer was added to the fractionation buffer, tracer was not added to the wash buffer. The supernatant (the ‘wash’ fraction) was quenched by transferal to a new 1.5 ml tube containing 0.125 ml of 50% (w/v) TCA in water to make a final concentration of 10% TCA. Residual wash was removed from the nuclear pellet, which was quenched by the resuspension in 1 ml 10% w/v TCA in water. Samples were stored at-80 before thawing on ice for processing or directly processed.

### 4.5 Whole cell extraction

For direct 10% TCA extraction: Media was poured from dishes into a waste container and cells were placed on ice at a 45° angle and residual media aspirated completely. Dishes were laid flat on ice and 1 ml 10% TCA in water added, cells were scraped into the TCA and transferred to 1.5 ml tubes kept on ice.

For 80:20 methanol:water extraction: Media was poured from dishes into a waste container and cells were placed on ice at a 45° angle and residual media aspirated completely. Aspirated dishes were laid flat on dry ice and 1 ml of −80 C 80:20 HPLC grade (Optima) methanol:water added, in which cells were scraped and transferred to 1.5 ml tubes kept on ice.

For PBS extraction: Cells were placed at a 45° angle on ice and medium was aspirated, cells were washed with 4 ml ice-cold PBS and aspirated completely. Before being scraped into 0.5 ml of ice-cold deionized water and transferred to a 1.5 ml Eppendorf tube containing 0.5 ml 4 °C 20% TCA in water.

For DMEM extraction: cells were placed flat on ice and scraped directly into the pre-incubation medium, transferred to 15 ml tubes (kept on ice) and spun down at 900 × g at 4 °C for 5 min. Media was removed by aspiration and the cell pellet resuspended in 1 ml 4 °C 10% TCA in water.

### 4.6 Acyl-CoA sample processing

Cell and fraction samples in 10% (w/v) trichloroacetic acid (Sigma cat. #T6399) in water were sonicated for 12 × 0.5 sec pulses, protein was pelleted by centrifugation at 17,000 × g from 10 min at 4 °C. The supernatant was purified by solid-phase extraction using Oasis HLB 1cc (30 mg) SPE columns (Waters). Columns were washed with 1 mL methanol, equilibrated with 1 mL water, loaded with supernatant, desalted with 1 mL water, and eluted with 1 mL methanol containing 25 mM ammonium acetate. The purified extracts were evaporated to dryness under nitrogen then resuspended in 55 μl 5% (w/v) 5-sulfosalicylic acid in water.

### 4.7 Acyl-CoA isotopologue analysis

Acyl-CoAs were measured by liquid chromatography-high resolution mass spectrometry. Briefly, 5-10 µl of purified samples in 5% SSA were analyzed by injection of an Ultimate 3000 HPLC coupled to a Q Exactive Plus (Thermo Scientific) mass spectrometer in positive ESI mode using the settings described previously[15]. This method was modified for the dual-labeling experiment by expansion of the isolation window to capture all possible isotopologues of succinyl-CoA and acetyl-CoA. Quantification of acyl-CoAs was via their MS2 fragments and the targeted masses used for isotopologue analysis are indicated in **Supp. Table 1**. Data were integrated using Tracefinder (Thermo Scientific) software and isotopic enrichment was calculated by normalization to unlabeled control samples using the FluxFix calculator[27].

### 4.8 TCA intermediate sample processing

HepG2 cells were grown in 10 cm dishes and pre-labeled as described above. At harvest, dishes were placed on ice, medium was aspirated thoroughly, and cells were immediately scraped into 1 ml/dish 80:20 Methanol:water (HPLC-grade, Optima) pre-chilled to −80 °C. Samples were transferred to 1.5 ml tubes and pulse-sonicated for 12 0.5 sec pulses with a probe tip sonicator and centrifuged at 17,000 rcf at 4 °C for 10 min. The supernatant was transferred to glass tubes and dried under nitrogen. Samples were resuspended in 100 μl of 5% w/v sulfosalicylic acid in water.

### 4.9 TCA intermediate isotopologue analysis

Isotopologue enrichment analysis of polar metabolites was performed using the ion-pairing reversed-phase ultra-high performance liquid chromatography method modified from Guo et al[28]. 5 μl was injected for analysis on Q Exactive HF instrument with SIM settings based on the unlabeled mass of each targeted metabolite with mass windows and offset accommodating all potential isotopologues. Masses used for isotopologue analysis are indicated in **Supp. Table 2**. Data from the MS1 peaks were integrated using XCalibur Quan browser (Thermo v2.3) software and isotopic enrichment was calculated by normalization to unlabeled control samples using the FluxFix calculator[27].

### 4.10 Stable isotope labeled substrates

^15^N_2_ ^13^C_5_-glutamine (Cambridge Isotope Laboratories CNLM-1275-H-PK), ^13^C_5_ -α-ketoglutarate (Cambridge Isotope Laboratories CNLM-2411-PK), ^13^C_2_-acetate (Cambridge Isotope Laboratories CNLM-440-5), ^13^C_6_-glucose (Cambridge Isotope Laboratories CLM-1396-1), Vitamin B5-[^13^C_6_ ^15^N_2_] (calcium pantothenate-[^13^C_6_ ^15^N_2_]) (Isosciences catalogue #5065).

### 4.11 Western blotting

Western blotting was performed using mini gel tank system (Life Biotechnologies) with 4-12% gradient Bis-Tris gels (NuPage, Invitrogen cat. #NP0335) and 0.45 µm pore size nitrocellulose membranes (BioRad cat. #1620115) with antibody incubations according to the manufacturers instructions. An Odyssey CLx imaging system with Image Studio v2.0.38 software (LI-COR Biosciences) was used to acquire images which were exported as TIFF files and cropped and arranged using Adobe Illustrator software v23.0.1 The following antibodies were used in this study; α-tubulin (Sigma, cat. #T16199), ACSS2 (Cell Signaling Technology, cat. #3658, clone: D19C6, lot: 2), ACLY (Proteintech, cat. #15421-1-AP, lot: 00040639), histone H3 (Abcam cat. #ab1791), histone H4 (Millipore cat. #05-858), FASN (Cell Signaling Technology, cat. #3189, lot: 2), citrate synthase (Cell Signaling Technology, cat. #14309, clone: D7V8B, lot: 1), LaminA/C (Cell Signaling Technology, cat. #2032), DNMT1 (Active Motif cat. # 39905 Lot# 12914002), LAMP1 (Cell Signaling Technology, cat. #3243 clone C54H11, lot 4), catalase (Cell Signaling Technologies, cat. #14097, clone D5N7V, lot 1).

### 4.12 Statistical analyses

Data presented are shown either of mean ± standard deviation or, for curve fits, mean ± 95% confidence intervals. Graphpad Prism software (v.8) was used for graphing and statistical analysis. One-phase association function was used to fit non-linear curves and calculate kinetic parameters including half-times. For comparison between two groups, datasets were analyzed by two-tailed Student’s t-test with Welch’s correction and statistical significance defined as p < 0.05 (*), p < 0.01 (**), p < 0.001 (***), p < 0.0001 (****).

## Acknowledgements

This work was funded by NIGMS grant R01GM132261 and NICHHD grant R03HD092630 to NWS and NCI grant R01CA174761 to KEW. ST was funded by American Diabetes Association post-doctoral fellowship #1-18-PDF-144. KH and JB-S were supported by the Austrian Science Fund grants FWF W1226 and FWF P27108.

## Author contributions

ST, NWS and KEW conceptualized the study and designed experiments. ST prepared figures and wrote the manuscript. NWS and KEW edited the manuscript. ST performed the majority of the experiments and data analysis. JL and KH performed experiments and analysis. MD, HJ, JS, EVK, AB and PX performed metabolite extraction and analysis. JB-S supported KH and provided useful discussion. All authors read and provided feedback on manuscript and figures.

## Data availability

All data relevant to this study are presented in the figures.

## Code availability

This study does not use any custom code that has not been published or licensed elsewhere.

---

**Supplemental Figure 1:**
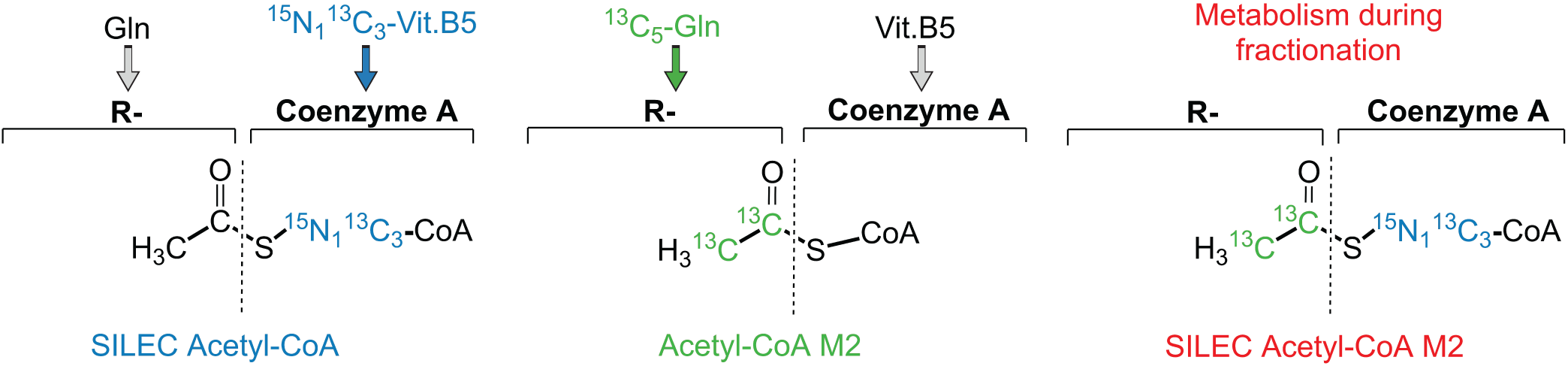
Dual labelling strategy reveals metabolism during fractionation. ^15^N_1_ ^13^C_3_-vitamin B5 is incorporated into the Coenzyme A backbone of acyl-CoA molecules (e.g. SILEC acetyl-CoA) whilst ^15^N_2_ ^13^C_5_-glutamine labels acyl (R-group) carbons (e.g. acetyl-CoA M4). Generation of doubly labelled molecules (e.g. SILEC succinyl-CoA M4) after the labelled cells are combined indicates post-harvest metabolism during fractionation.

**Supplemental Figure 2:**
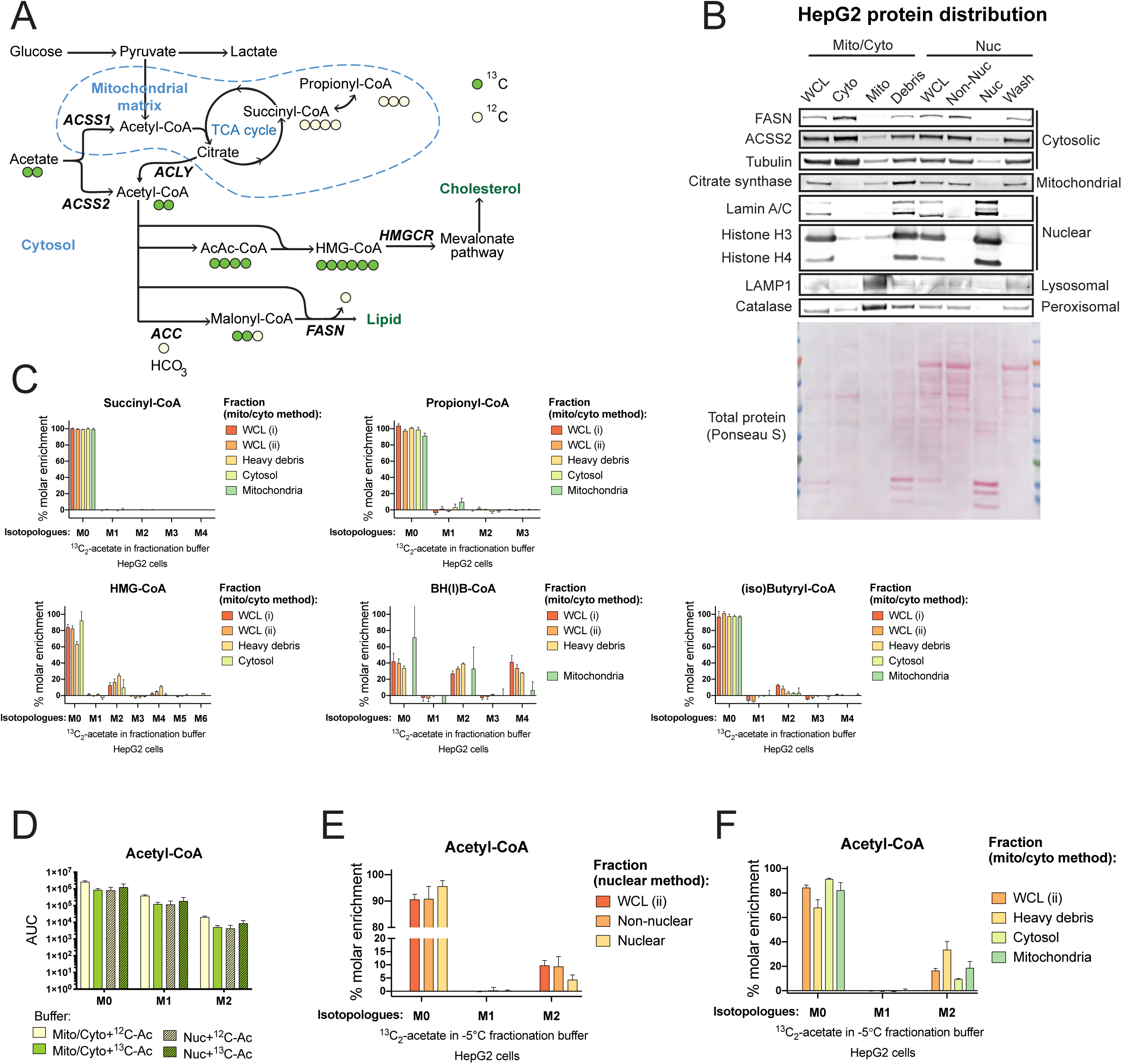
Substrate use during fractionation follows defined metabolic pathways. **A**) Schematic representation of acetate metabolism in HepG2 cells. **B**) Western blotting was used to confirm appropriate protein distribution in sub-cellular fractions from HepG2 cells with equal protein loading. FASN (fatty acid synthase), ACSS2 (Acyl-CoA Synthetase Short Chain Family Member 2), LAMP1 (lysosomal associated membrane protein 1), WCL (whole cell lysate), Cyto (cytosol), mito (mitochondria). **C**) Isotopologue enrichment of short chain acyl-CoA species following addition of 0.1 mM ^13^C_2_-acetate to the fractionation buffer during mito/cyto fractionation procedure in HepG2 cells. 3-hydroxymethylglutaryl-CoA (HMG-CoA), β-hydroxy(iso)butyryl-CoA (BH(I)B-CoA). Isobutyryl-CoA and butyryl-CoA, and β-hydroxyisobutyryl-CoA and β-hydroxybutyryl-CoA were not differentiated by mass spectrometry analysis. **D**) Raw area under the curve (AUC) data from isotopologue peaks of acetyl-CoA in de-proteinated HepG2 cell extracts resuspended in fractionation buffer for mito/cyto or nuclear protocols containing 0.1 mM ^13^C_2_-acetate or unlabeled acetate and incubated on ice for 1 h before extraction. **E-F**) HepG2 cells were harvested at −5 °C in fractionation buffer containing 0.1 mM ^13^C_2_-acetate and % molar enrichment of acetyl-CoA isotopologues was calculated by normalization to control samples with unlabeled acetate in the fractionation buffer for mito/cyto and nuclear fractionation protocols. Mean of n=3 replicates from representative experiments is shown with error bars indicating standard deviation.

**Supplemental Figure 3:**
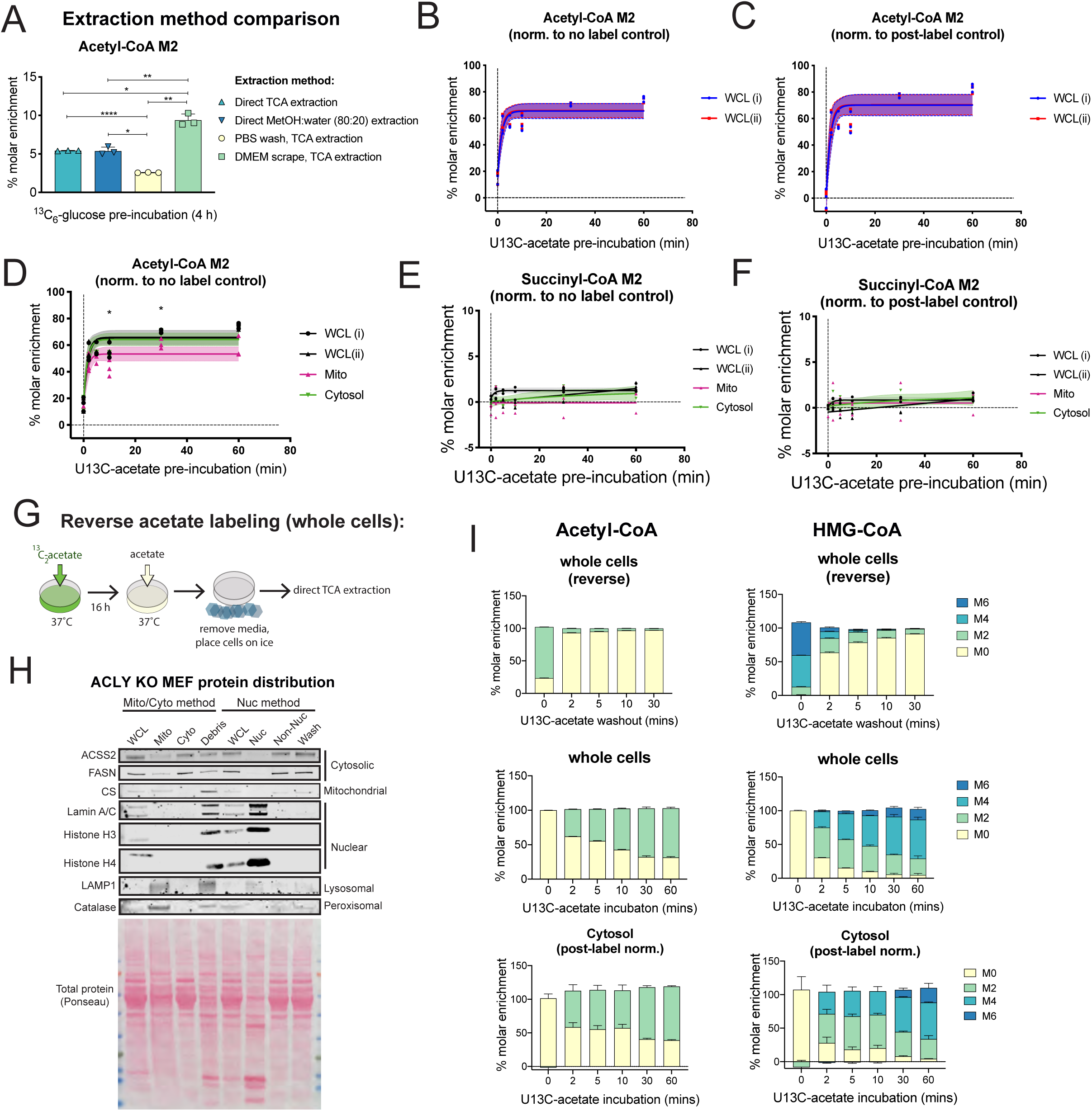
Post-labeling correction resolves mitochondrial and cytosolic acetyl-CoA metabolism. **A**) Comparison of incorporation of glucose into acetyl-CoA in ACLY KO MEFs pre-incubated in 5 mM ^13^C_6_-glucose in the absence of acetate for 4 h before harvest of whole cells using 4 different techniques. **B-H**) ACLY KO MEF cells were pre-labeled by incubation with 0.1 mM ^13^C_2_-acetate over a time-course. **B**) Comparison of absolute labeling in WCL i and WCLI ii post-labeled samples by normalization to no label control. **D**) Comparison of corrected WCL i and WCLI ii post-labeled data when normalized to post-label control. **D-E)** Absolute isotopologue enrichment of mito/cyto fractionation post-labeled data when normalized to no label control for acetyl-CoA M2 and succinyl-CoA M2. **F**) Succinyl-CoA M2 post-labeled data when normalized to post-label control. **G**) Schematic representation of reverse labeling scheme with 16 h pre-incubation in 0.1 mM ^13^C_2_-acetate switched to unlabeled acetate over a time-coursse before direct extraction of whole cells in trichloroacetic acid (TCA). **H**) Protein distribution analysis by Western blot in sub-cellular fractions from ACLY KO MEFs with equal protein loading. CS (citrate synthase), ACSS2 (Acyl-CoA Synthetase Short Chain Family Member 2). **I**) Comparison of acetyl-CoA and HMG-CoA isotopologue enrichment in whole cells with direct extraction (reverse and forward labeling) and cytosol isolated by sub-cellular fractionation with post-labeling normalization with the addition 5 mM each of ^13^C_2_ -acetate and unlabeled acetate in the fractionation buffer (forward labeling). A representative expreriment for each paradigm is shown with n= 3 distinct replicates for each experiment. Error bars represent standard deviation and induvidual replicates are not displayed for the sake of graphical clarity. **A-F**) Data from representative experiments is shown. Shading indicates 95% confidence bands and symbols represent individual replicate values (n=3-4). For comparison between two groups, datasets were analyzed by student’s t-test with statistical significance defined as p < 0.05 (*), p < 0.01 (**), p < 0.001 (***), p < 0.0001 (****).

**Supplemental Figure 4:**
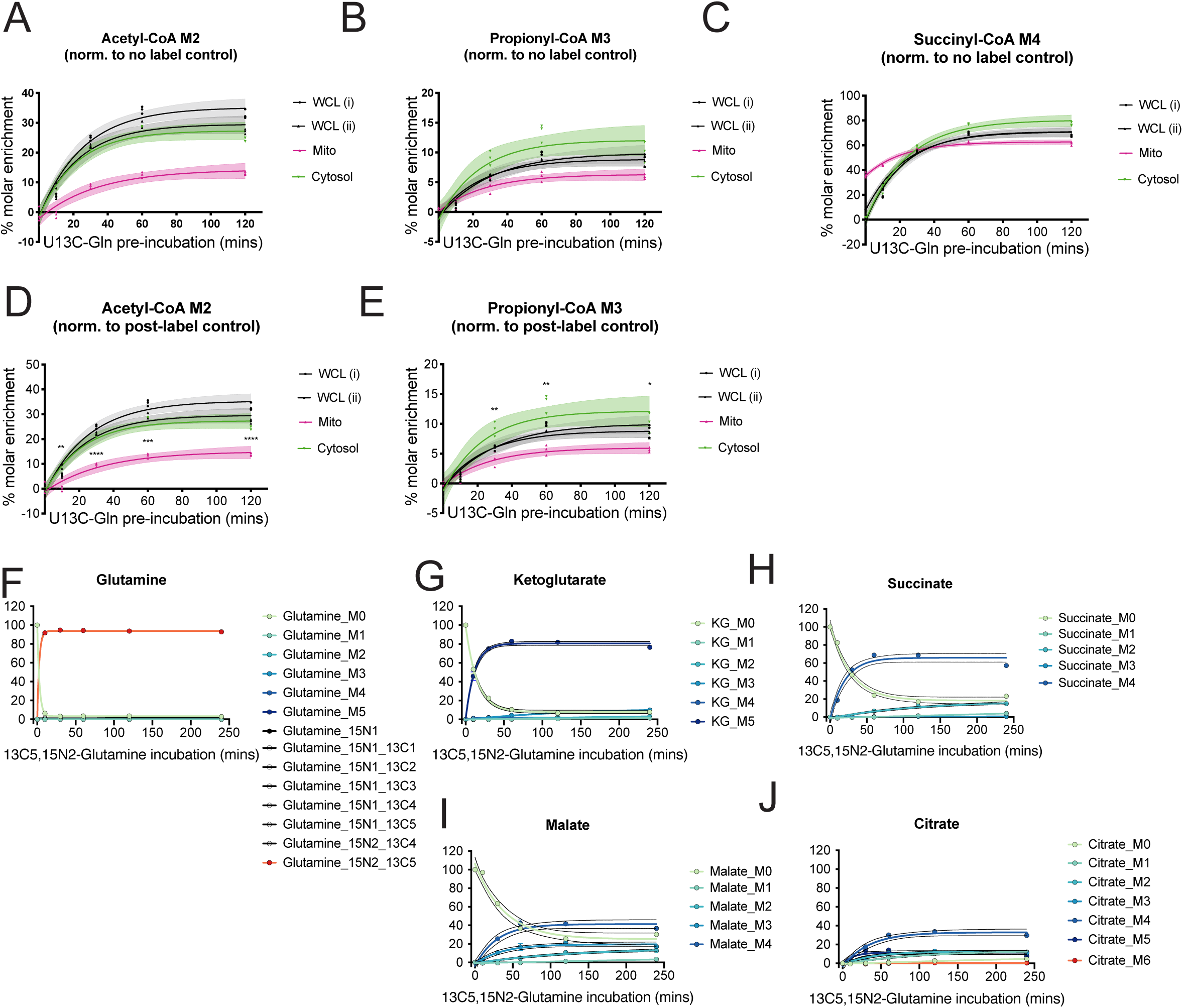
Kinetics of glutamine metabolism in the mitochondria are transformed by post-label correction. HepG2 cells were pre-labeled over a time-course with 2 mM ^15^N_2_ ^13^C_5_-glutamine. Absolute isotopologue enrichment of mito/cyto fractionation post-labeled data when normalized to no label control for acetyl-CoA M2 (**A**), propionyl-CoA M3 (**B**) and succinyl-CoA M4 (**C**). Acetyl-CoA M2 (**D**) and propionyl-CoA M3 (**E**) post-labeled data when corrected by normalization to post-labeled control. **F-J**) Isotopologue analysis of tricarboxylic acid (TCA) cycle intermediates after direct extraction from whole cells. Data from representative experiments is shown. Shading indicates 95% confidence bands and symbols represent individual replicate values (n=3). For comparison between two groups, datasets were analyzed by student’s t-test with statistical significance defined as p < 0.05 (*), p < 0.01 (**), p < 0.001 (***), p < 0.0001 (****).

