## Supplemental Table 1 for "Subcellular metabolic pathway kinetics are revealed by correcting for artifactual post harvest metabolism"

**Table 1: Acyl-CoA Mass List**

| Isotopologue | Precursor Formula [M] | Precursor Ion [M+H]+ | Product Ion [M+H-507]+ |
|---|---|---|---|
| Succinyl-CoA | C25H40N7O19P3S | 868.1385 | 361.1428 |
| Succinyl-CoA-M1 | [13]C1C24H40N7O19P3S | 869.1419 | 362.1461 |
| Succinyl-CoA-M2 | [13]C2C23H40N7O19P3S | 870.1452 | 363.1495 |
| Succinyl-CoA-M3 | [13]C3C22H40N7O19P3S | 871.1486 | 364.1528 |
| Succinyl-CoA-M4 | [13]C4C21H40N7O19P3S | 872.1520 | 365.1562 |
| Succinyl-CoA-M5 | [13]C5C20H40N7O19P3S | 873.1553 | 366.1596 |
| Succinyl-CoA-M6 | [13]C6C19H40N7O19P3S | 874.1587 | 367.1629 |
| Succinyl-CoA-M7 | [13]C7C18H40N7O19P3S | 875.1620 | 368.1663 |
| Acetyl-CoA | C23H38N7O17P3S | 810.1331 | 303.1373 |
| Acetyl-CoA-M1 | [13]C1C22H38N7O17P3S | 811.1364 | 304.1407 |
| Acetyl-CoA-M2 | [13]C2C21H38N7O17P3S | 812.1398 | 305.1440 |
| Acetyl-CoA-M3 | [13]C3C20H38N7O17P3S | 813.1431 | 306.1474 |
| Acetyl-CoA-M4 | [13]C4C19H38N7O17P3S | 814.1465 | 307.1507 |
| Acetyl-CoA-M5 | [13]C5C18H38N7O17P3S | 815.1498 | 308.1541 |
| Propionyl-CoA | C24H40N7O17P3S | 824.1487 | 317.1530 |
| Propionyl-CoA-M1 | [13]C1C23H40N7O17P3S | 825.1521 | 318.1563 |
| Propionyl-CoA-M2 | [13]C2C22H40N7O17P3S | 826.1554 | 319.1488 |
| Propionyl-CoA-M3 | [13]C3C20H40N7O17P3S | 827.1588 | 320.1521 |
| Propionyl-CoA-M4 | [13]C4C19H40N7O17P3S | 828.1621 | 321.1555 |
| Propionyl-CoA-M5 | [13]C5C18H40N7O17P3S | 829.1655 | 322.1564 |
| 3HMG-CoA* | C27H44N7O20P3S | 912.1647 | 405.1690 |
| 3HMG-CoA-M1 | [13]C1C26H44N7O20P3S | 913.1681 | 406.1724 |
| 3HMG-CoA-M2 | [13]C2C25H44N7O20P3S | 914.1715 | 407.1757 |
| 3HMG-CoA-M3 | [13]C3C24H44N7O20P3S | 915.1748 | 408.1791 |
| 3HMG-CoA-M4 | [13]C4C23H44N7O20P3S | 916.1782 | 409.1824 |
| 3HMG-CoA-M5 | [13]C5C22H44N7O20P3S | 917.1815 | 410.1858 |
| 3HMG-CoA-M6 | [13]C6C21H44N7O20P3S | 918.1849 | 411.1891 |
| 3HMG-CoA-M7 | [13]C7C20H44N7O20P3S | 919.1882 | 412.1925 |
| 3HMG-CoA-M8 | [13]C8C19H44N7O20P3S | 920.1916 | 413.1958 |
| 3HMG-CoA-M9 | [13]C9C18H44N7O20P3S | 921.1949 | 414.1992 |
| 3H(I)B-CoA** | C25H42N7O18P3S | 854.1593 | 347.1635 |
| 3H(I)B -CoA-M1 | [13]C1C24H42N7O18P3S | 855.1626 | 348.1669 |
| 3H(I)B -CoA-M2 | [13]C2C23H42N7O18P3S | 856.1660 | 349.1702 |
| 3H(I)B -CoA-M3 | [13]C3C22H42N7O18P3S | 857.1693 | 350.1736 |
| 3H(I)B -CoA-M4 | [13]C4C21H42N7O18P3S | 858.1727 | 351.1769 |
| 3H(I)B -CoA-M5 | [13]C5C20H42N7O18P3S | 859.1760 | 352.1803 |
| 3H(I)B -CoA-M6 | [13]C6C19H42N7O18P3S | 860.1794 | 353.1836 |
| 3H(I)B -CoA-M7 | [13]C7C18H42N7O18P3S | 861.1827 | 354.1870 |
| [Iso]Butyryl-CoA | C25H42N7O17P3S | 838.1644 | 331.1686 |
| [Iso]Butyryl-CoA-M1 | [13]C1C24H42N7O17P3S | 839.1677 | 332.1720 |
| [Iso]Butyryl-CoA-M2 | [13]C2C23H42N7O17P3S | 840.1711 | 333.1753 |
| [Iso]Butyryl-CoA-M3 | [13]C3C22H42N7O17P3S | 841.1744 | 334.1787 |
| [Iso]Butyryl-CoA-M4 | [13]C4C21H42N7O17P3S | 842.1778 | 335.1820 |
| [Iso]Butyryl-CoA-M5 | [13]C5C20H42N7O17P3S | 843.1811 | 336.1854 |
| [Iso]Butyryl-CoA-M6 | [13]C6C19H42N7O17P3S | 844.1845 | 337.1887 |
| [Iso]Butyryl-CoA-M7 | [13]C7C18H42N7O17P3S | 845.1878 | 338.1921 |
| Acetyl-CoA SILEC | [13]C3C20H38[15]N1N6O17P3S | 814.1401 | 307.1444 |
| Acetyl-CoA SILEC-M1 | [13]C4C19H38[15]N1N6O17P3S | 815.1435 | 308.1478 |
| Acetyl-CoA SILEC-M2 | [13]C5C18H38[15]N1N6O17P3S | 816.1469 | 309.1511 |
| Acetyl-CoA SILEC-M3 | [13]C6C17H38[15]N1N6O17P3S | 817.1502 | 310.1545 |
| Acetyl-CoA SILEC-M4 | [13]C7C16H38[15]N1N6O17P3S | 818.1536 | 311.1578 |
| Acetyl-CoA SILEC-M5 | [13]C8C15H38[15]N1N6O17P3S | 819.1569 | 312.1612 |
| Succinyl-CoA SILEC | [13]C3C22H40[15]N1N6O19P3S | 872.1456 | 365.1499 |
| Succinyl-CoA SILEC-M1 | [13]C4C21H40[15]N1N6O19P3S | 873.1490 | 366.1533 |
| Succinyl-CoA SILEC-M2 | [13]C5C20H40[15]N1N6O19P3S | 874.1523 | 367.1566 |
| Succinyl-CoA SILEC-M3 | [13]C6C19H40[15]N1N6O19P3S | 875.1557 | 368.1560 |
| Succinyl-CoA SILEC-M4 | [13]C7C18H40[15]N1N6O19P3S | 876.1591 | 369.1633 |
| Succinyl-CoA SILEC-M5 | [13]C8C17H40[15]N1N6O19P3S | 877.1624 | 370.1666 |
| Succinyl-CoA SILEC-M6 | [13]C9C16H40[15]N1N6O19P3S | 878.1658 | 371.1700 |
| Succinyl-CoA SILEC-M7 | [13]C10C15H40[15]N1N6O19P3S | 879.1691 | 372.1734 |

*HMG-CoA (3-hydroxymethylglutaryl-CoA), **3H(I)B-CoA (3-hydroxybutyrate-CoA/3-hydroxyisobutyrate-CoA)

**Table 2: TCA Cycle Mass List**

| Isotopologue | Formula [M] | Precursor Ion [M-H]- |
|---|---|---|
| Glutamine | C5H10N2O3 | 145.0608 |
| [13]C1-glutamine | [13]C1C4H10N2O3 | 146.0641 |
| [13]C2-glutamine | [13]C2C3H10N2O3 | 147.0675 |
| [13]C3-glutamine | [13]C3C2H10N2O3 | 148.0708 |
| [13]C4-glutamine | [13]C4C1H10N2O3 | 149.0742 |
| [13]C5-glutamine | [13]C5H10N2O3 | 150.0775 |
| [15]N1-glutamine | C5H10[15]N1N1O3 | 146.0589 |
| [13]C1[15]N1-glutamine | [13]C1C4H10[15]N1N1O3 | 147.0623 |
| [13]C2[15]N1-glutamine | [13]C2C3H10[15]N1N1O3 | 148.0656 |
| [13]C3[15]N1-glutamine | [13]C3C2H10[15]N1N1O3 | 149.0690 |
| [13]C4[15]N1-glutamine | [13]C4C1H10[15]N1N1O3 | 150.0723 |
| [13]C5[15]N1-glutamine | [13]C5H10[15]N1N1O3 | 151.0757 |
| [15]N2-glutamine | C5H10[15]N2O3 | 147.0559 |
| [13]C1[15]N2-glutamine | [13]C1C4H10[15]N2O3 | 148.0593 |
| [13]C2[15]N2-glutamine | [13]C2C3H10[15]N2O3 | 149.0626 |
| [13]C3[15]N2-glutamine | [13]C3C2H10[15]N2O3 | 150.0660 |
| [13]C4[15]N2-glutamine | [13]C4C1H10[15]N2O3 | 151.0694 |
| [13]C5[15]N2-glutamine | [13]C5H10[15]N2O3 | 152.0727 |
| $\alpha$-Ketoglutarate | C5H6O5 | 145.0142 |
| [13]C1-$\alpha$-Ketoglutarate | [13]C1C4H6O5 | 146.0176 |
| [13]C2-$\alpha$-Ketoglutarate | [13]C2C3H6O5 | 147.0185 |
| [13]C3-$\alpha$-Ketoglutarate | [13]C3C2H6O5 | 148.0218 |
| [13]C4-$\alpha$-Ketoglutarate | [13]C4C1H6O5 | 149.0227 |
| [13]C5-$\alpha$-Ketoglutarate | [13]C5H6O5 | 150.0261 |
| Succinate | C4H5O4 | 117.0193 |
| [13]C1-succinate | [13]C1C3H5O4 | 118.0227 |
| [13]C2-succinate | [13]C2C2H5O4 | 119.0236 |
| [13]C3-succinate | [13]C3C1H5O4 | 120.0269 |
| [13]C4-succinate | [13]C4H5O4 | 121.0278 |
| Malate | C4H6O5 | 133.0131 |
| [13]C1-malate | [13]C1C3H6O5 | 134.0165 |
| [13]C2-malate | [13]C2C2H6O5 | 135.0199 |
| [13]C3-malate | [13]C3C1H6O5 | 136.0232 |
| [13]C4-malate | [13]C4H6O5 | 137.0266 |
| Citrate | C6H8O7 | 191.0197 |
| [13]C1-citrate | [13]C1C5H8O7 | 192.0231 |
| [13]C2-citrate | [13]C2C4H8O7 | 193.0240 |
| [13]C3-citrate | [13]C3C3H8O7 | 194.0273 |
| [13]C4-citrate | [13]C4C2H8O7 | 195.0331 |
| [13]C5-citrate | [13]C5C1H8O7 | 196.0365 |
| [13]C6-citrate | [13]C6H8O7 | 197.0374 |
